# Pathogenic LRRK2 mutations cause loss of primary cilia and Neurturin in striatal Parvalbumin interneurons

**DOI:** 10.1101/2024.06.17.599289

**Authors:** Yu-En Lin, Ebsy Jaimon, Francesca Tonelli, Suzanne R. Pfeffer

**Author notes:** To whom correspondence should be addressed: Suzanne Pfeffer, Department of Biochemistry, 279 Campus Drive Beckman B400, Stanford University School of Medicine, Stanford, CA 94305-5307.

## Abstract

Parkinson’s disease-associated, activating mutations in LRRK2 kinase block primary cilia formation in cell culture and in specific cell types in the brain. In the striatum that is important for movement control, about half of astrocytes and cholinergic interneurons, but not the predominant medium spiny neurons, lose their primary cilia. Here we show that Parvalbumin interneurons that are inhibitory regulators of movement also lose primary cilia. Without cilia, these neurons are not able to respond to Sonic hedgehog signals that normally induce the expression of Patched protein, and their numbers decrease. In addition, glial cell line-derived neurotrophic factor-related Neurturin expression is significantly decreased. These experiments highlight the importance of Parvalbumin neurons in cilia-dependent, neuroprotective signaling pathways and show that LRRK2 activation decreases Neurturin production, resulting in less neuroprotection for dopamine neurons.

**Summary:** Parvalbumin interneurons in the dorsal striatum lose primary cilia in mice harboring Parkinson’s-associated, activating mutations in LRRK2 kinase, resulting in loss of Hedgehog signaling and decreased production of neuroprotective, Glial cell line-derived neurotrophic factor-related Neurturin to support dopamine neurons.

## Introduction

Parkinson’s disease is characterized by the specific loss of dopamine-producing neurons that project into a region of the brain known as the dorsal striatum (Morris et al., 2024). The release of dopamine in the striatum enhances motor functions and reward signaling; additionally, dopamine plays a role in reinforcement learning (Balleine et al., 2007; Jurado-Parras et al., 2020). Imbalances in dopamine levels in the striatum are central to Parkinson’s disease.

The primary cells in the striatum that respond to dopamine are known as medium spiny neurons (Balleine et al., 2007; Jurado-Parras et al., 2020; Kreitzer and Malenka, 2008). The dorsal striatum also contains multiple populations of interneurons, each comprising only a small fraction of the total cell population (Chang and Kita, 1992; Munoz-Manchado et al., 2018; Stanley et al., 2020). Among the most well-characterized striatal interneurons are the Parvalbumin (PV) and cholinergic cell types (Gritton et al., 2019; Tanimura et al., 2018). PV neurons release GABA and inhibit the activity of nearby medium spiny neurons, aiding in motor control (Gittis et al., 2011; Koos and Tepper, 1999; Nahar et al., 2021; Tepper et al., 2018). On the other hand, cholinergic interneurons release acetylcholine to modulate the activity of medium spiny neurons and are also involved in motor output control. Recent studies suggest that PV neurons fine-tune the activation of medium spiny neuron networks essential for movement execution, whereas cholinergic interneurons help synchronize activity within medium spiny neuron networks, signaling the end of movement.

We study an inherited form of Parkinson’s that is caused by activating mutations in the Leucine Rich Repeat Kinase 2 (LRRK2) (Alessi and Sammler, 2018). LRRK2 kinase phosphorylates a subset of Rab GTPases (Steger et al., 2017; Steger et al., 2016) and complexes of phosphoRab10 bound to RILPL1 protein block the formation of primary cilia in multiple cultured cell lines (Dhekne et al., 2018; Steger et al., 2017) but only in certain cell types in the brain (Dhekne et al., 2018). In cell culture, cilia blockade occurs at the earliest steps of ciliogenesis: hyperactive LRRK2 blocks the recruitment of tau tubulin kinase 2 to the mother centriole to trigger release of CP110, thus blocking cilia formation (Sobu et al., 2021).

We have shown that in the dorsal striatum in multiple LRRK2 pathway mouse models and in humans harboring LRRK2 mutations, rare cholinergic interneurons and astrocytes lose cilia but the much more abundant, surrounding medium spiny neurons do not (Dhekne et al., 2018; Khan et al., 2024; Khan et al., 2021). Loss of cilia correlates with an inability to sense and respond to Sonic hedgehog (Shh) signals (Dhekne et al., 2018) that require cilia for signal transduction (Corbit et al., 2005; Dhekne et al., 2018; Rohatgi et al., 2007). In the absence of Shh, or in the presence of pathogenic LRRK2, cholinergic interneurons decrease production of glial derived neurotrophic factor (GDNF) that normally supports dopamine neurons (Gonzalez-Reyes et al., 2012; Khan et al., 2024). In addition, Shh signaling is needed for cholinergic interneuron survival (Gonzalez-Reyes et al., 2012).

Here we report the consequences of LRRK2 mutation on PV neurons that are critical regulatory contributors to the nigrostriatal circuit. We show that like cholinergic interneurons, rare PV neurons also lose primary cilia and their ability to carry out Shh signaling; the consequence of PV neuron cilia loss is a major reduction of GDNF-related Neurturin (NRTN) expression, and a loss of cell numbers, decreasing neuroprotection for vulnerable dopamine neurons, and contributing to Parkinson’s disease.

## Results and Discussion

We analyzed the primary ciliation status of PV and Somatostatin interneurons in the mouse dorsal striatum of wild type and LRRK2 G2019S mice. As shown in Figure 1 (A, B), PV neurons were approximately 70% ciliated in wild type 5-month-old mice, as monitored using adenylate cyclase 3 (ACIII) antibodies to detect neuronal cilia. In contrast, PV neurons in the striatum of mice harboring the LRRK2 G2019S mutation showed a roughly 30% decrease in their overall ciliation status (Fig. 1A,B). In addition, the remaining cilia were ∼30% shorter which decreases their overall signaling capacity (Fig. 1D). In Somatostatin interneurons detected using anti-SST28 antibodies, loss of primary cilia was slightly less pronounced at ∼25% but still significant (Fig. 1A,C). In this study, we focused subsequent analysis on the PV neuron class.

**Figure 1.**
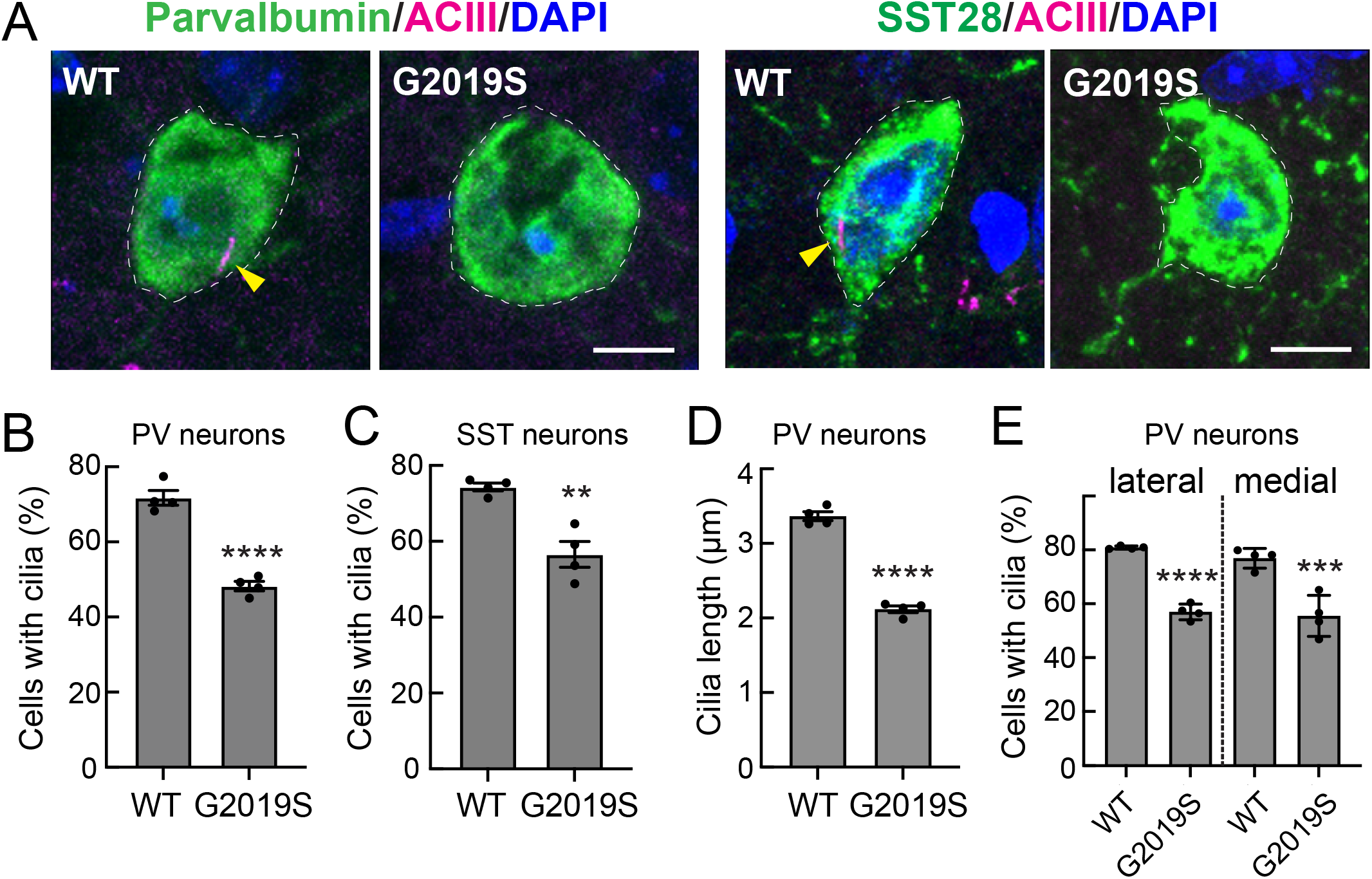
G2019S LRRK2 striatal parvalbumin and somatostatin interneurons have fewer cilia. (A) Example confocal immunofluorescence micrographs of sections of the dorsal striatum from 5-month-old wild-type (WT) or G2019S LRRK2 knock-in mice; scale bar, 5 μm. For every cell scored, care is taken to ensure that each cilium is associated with a specific cell body and emanates from near the nucleus. (A) Parvalbumin (PV) interneuron cell bodies were detected with anti-PV antibody (green, dashed white outline); primary cilia were detected with anti-Adenylate cyclase antibody (ACIII, magenta, highlighted by yellow arrowheads); nuclei were detected by DAPI (blue) staining. Somatostatin interneurons were detected with anti-Somatostatin-28 antibody (SST28, green, dashed white outline); primary cilia and nuclei were detected as in (A). (B) Percentage of PV^+^ neurons containing a cilium. (C) Percentage of SST^+^ neurons containing a cilium. (D) Quantification of the cilia length of PV neurons. Significance was determined by unpaired *t*-test; (B) ****, p < 0.0001; (C) **, p = 0.0025. (D) ****, p < 0.0001. Values represent the mean ± SEM from individual brains, analyzing 4 brains per group, 2-3 sections per mouse, and >40 neurons per mouse. (E) Percentage of PV neurons containing a cilium in the dorsolateral or dorsomedial striatum for wild type and G2019S mice. Significance was determined by one-way ANOVA with Tukey’s test; ****, p <0.0001; ***, p = 0.0001. Values represent the data (mean ± SEM) from individual brains, analyzing 4 brains per group, 2-3 sections per mouse, and >30 neurons per mouse.

PV interneurons are heterogenous: those in the dorso-medial striatum have been reported to have increased excitability compared to those in the dorsolateral striatum; they also uniquely receive glutamatergic input from the cingulate cortex (Tepper et al., 2018). Nevertheless, we detected a similar loss of cilia in both dorsolateral and dorsomedial PV neuron classes (Fig. 1E).

### Cilia loss correlates with G2019S LRRK2 expression

To investigate the basis for cilia loss in PV neurons, we employed RNAscope™ fluorescence in situ hybridization to monitor the cell type-specific expression of various gene products. In these experiments, antibodies were used to detect PV neurons and their ciliation status; RNA hybridization probes were used to detect specific gene products in the identified cell types. Figure 2A shows examples of ciliated and non-ciliated cells from wild type or LRRK2 G2019S dorsal striatum (left column) and their corresponding content of LRRK2 mRNA (right column). Individual dots reflect amplified signals from single RNA transcripts. Quantitation of at least 100 ciliated or non-ciliated PV neurons from four mice showed that non-ciliated wild type and non-ciliated LRRK2 G2019S PV neurons express higher levels of LRRK2 enzyme than their ciliated counterparts (Figure 2B). Thus, within the class of PV cells, higher LRRK2 expression correlates directly with cilia loss.

**Figure 2.**
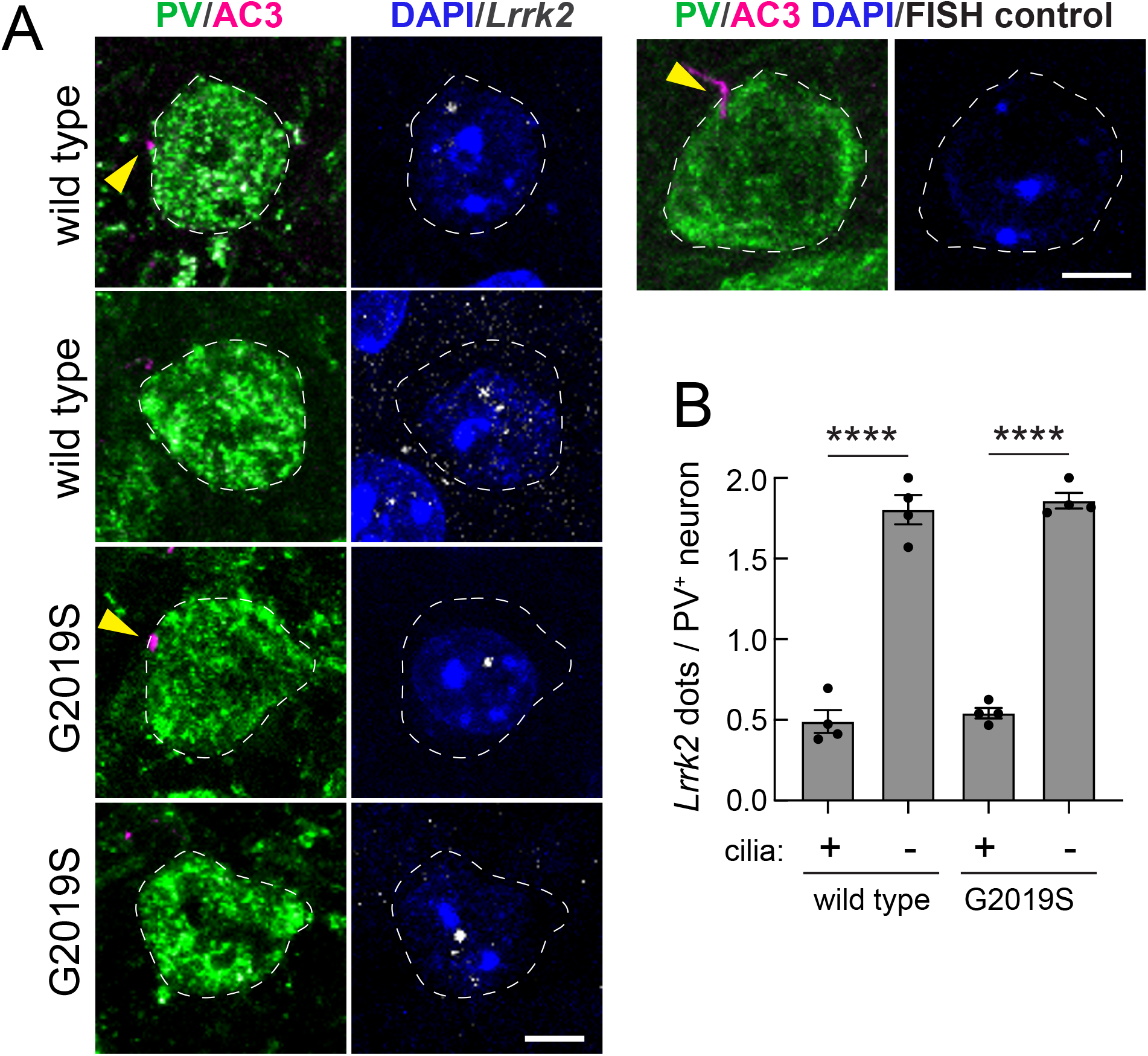
Loss of primary cilia in parvalbumin interneurons correlates with higher LRRK2 expression. (A) Example confocal immunofluorescence microscopy to identify PV neurons and their cilia as in Fig. 1A (left column), coupled with RNAscope in situ hybridization to detect LRRK2 transcripts in the same cells (right column). The images of a control neuron without an RNAscope probe are shown above panel B on the top right. Bar, 5 μm. (B) Quantitation of LRRK2 RNA dots per neuron as a function of ciliation status for wild type and G2019S mice. Significance was determined by one-way ANOVA with Tukey’s test; ****, p < 0.0001. Values represent the mean ± SEM from individual brains, analyzing 4 brains per group, 2-3 sections per mouse, and >25 neurons per mouse.

### Cilia loss correlates with loss of Sonic hedgehog signaling

Primary cilia are required to transduce the Shh signal (Corbit et al., 2005; Rohatgi et al., 2007). We thus predicted that cilia loss should lead to decreased expression of Shh target genes. Figure 3 shows that expression of Patched (Ptch1) RNA that encodes the Shh receptor was strongly decreased in mice harboring the hyperactive G2019S LRRK2 protein (Fig. 3B), to an extent predicted from a combination of cilia loss and ciliary length decrease (Fig. 1B, D). When the data were segregated based upon ciliation status, ciliated wild type PV neurons expressed the highest levels of Ptch1 RNA; much less was seen in non-ciliated PV neurons (Fig. 3C). A roughly 3-fold decrease in Ptch1 expression was seen even in the ciliated G2019S mutant cells, suggesting that the remaining, shorter cilia are less able to support Shh signaling. These data confirm the cilia dependence of Shh signaling in wild type striatal PV neurons and reveal a severe deficit in Shh signaling in these neurons in G2019S LRRK2 mice.

**Figure 3.**
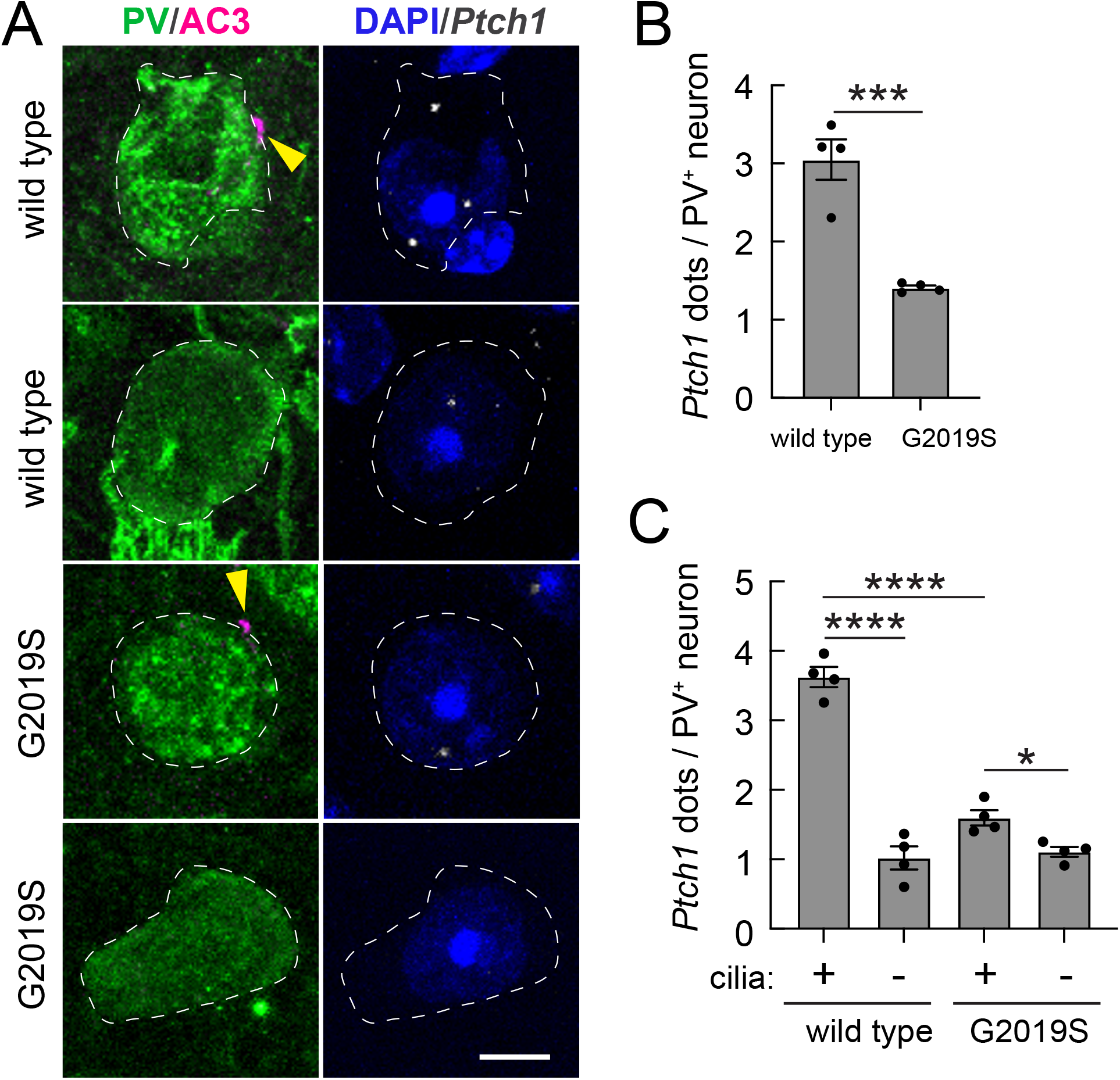
Decreased cilia-dependent PTCH1 expression in striatal G2019S LRRK2 parvalbumin interneurons. (A) Example confocal immunofluorescence microscopy to identify PV neurons and their cilia as in Fig. 1A (left column), coupled with RNAscope in situ hybridization to detect PTCH1 transcripts (right column) as in Figure 2. Bar, 5 μm. (B, C) Quantitation of PTCH1 RNA dots per neuron (B) or as a function of ciliation status (C) for wild type and G2019S mice. Significance was determined by unpaired *t*-test (B) and one-way ANOVA with Tukey’s test (C). (B) ***, p = 0.0007. (C) *, p = 0.0348; ****, p < 0.0001. Values represent the data (mean ± SEM) from individual brains, analyzing 4 brains per group, 2-3 sections per mouse, and >30 neurons per mouse.

### Loss of GDNF-related Neurturin production in LRRK2 G2019S PV neurons

Neurturin (NRTN) is a member of the glial cell line-derived neurotrophic factor (GDNF) family, which plays a critical role in the survival and differentiation of dopaminergic neurons (Kotzbauer et al., 1996). Within the striatum, NRTN is produced almost exclusively by PV neurons, with a small contribution from cholinergic neurons (Saunders et al., 2018). Because GDNF production is decreased in the absence of Shh ligand (Gonzalez-Reyes et al., 2012) and GDNF RNA is decreased in the absence of primary cilia (Khan et al., 2024), we assume that GDNF is either a direct, or indirect, Shh target gene. We thus explored whether PV neurons express less GDNF-related NRTN in conjunction with LRRK2 G2019S cilia-blockade and Shh signaling dysfunction.

Figure 4A shows the results of RNAscope™ analysis of NRTN transcripts. As predicted, we detected an almost 50% loss of NRTN RNA in LRRK2 G2019S striatal PV neurons (Fig. 4B), directly analogous to the extent of Shh signaling loss (Fig. 3B). When segregating the data according to ciliation status, we noted that in wild type brain, ciliated PV neurons express about three times more NRTN RNA than non-ciliated PV neurons. In contrast, similar to what we observed for PTCH1 RNA expression, even ciliated PV neurons in LRRK2 G2019S dorsal striatum showed decreased NRTN expression, consistent with a ciliary signaling defect. Altogether, these data show that LRRK2 G2019S interferes with PV neuron ciliogenesis, Shh signaling and production of the neuroprotective NRTN protein.

**Figure 4.**
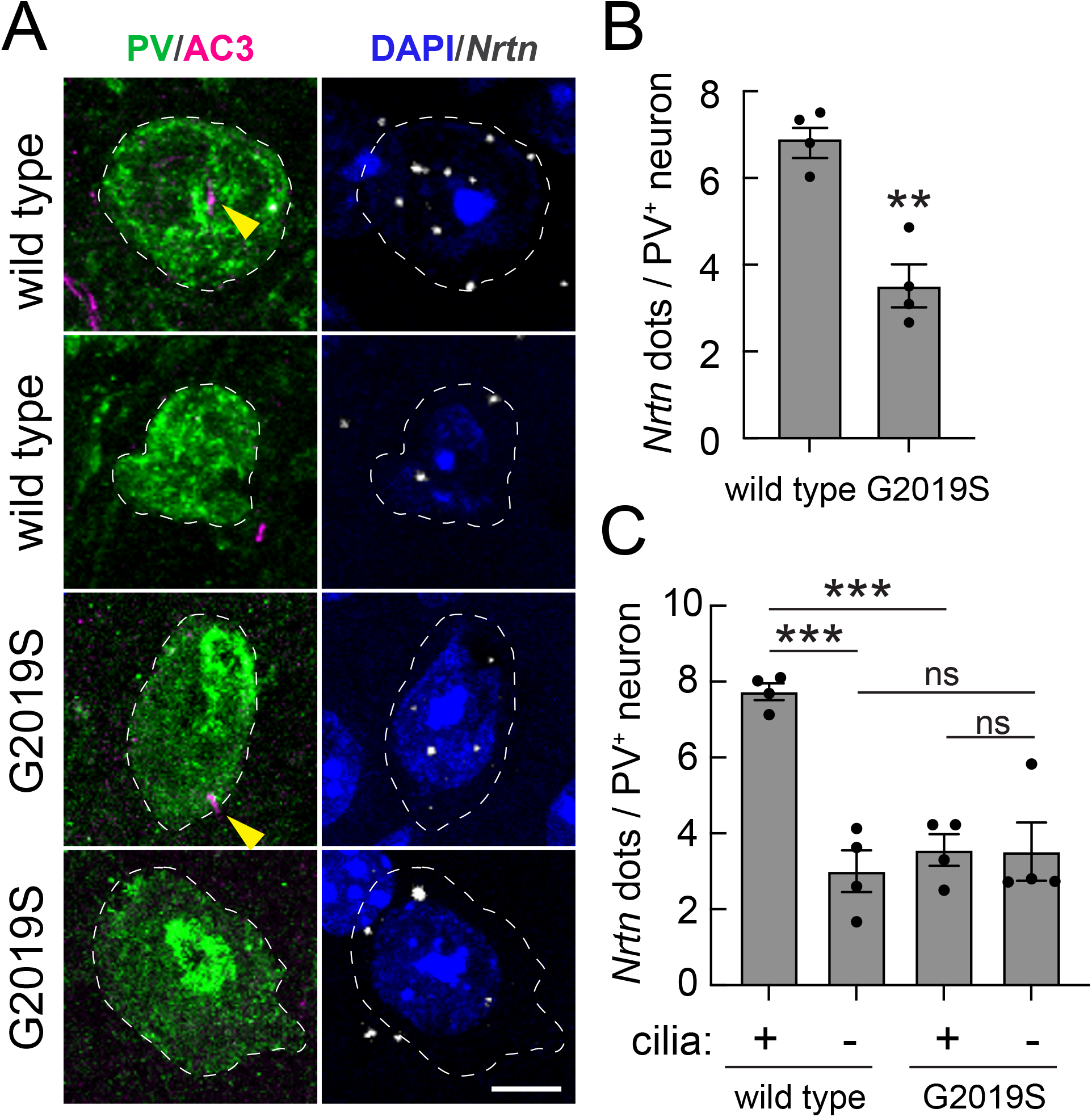
Decreased cilia-dependent Neurturin expression in striatal G2019S LRRK2 parvalbumin interneurons. (A) Confocal immunofluorescence microscopy to identify PV neurons and their cilia as in Fig. 1A (left column), coupled with RNAscope in situ hybridization to detect NRTN transcripts (right column) as in Figure 2. Bar, 5 μm. (B, C) Quantitation of Nrtn RNA dots per neuron (B) or as a function of ciliation status (C) for wild type and G2019S mice. Significance was determined by unpaired *t*-test (B) and one-way ANOVA with Tukey’s test (C). (B) **, p = 0.0011. (C) ***, p = 0.0002 between ciliated and nonciliated wild type mice; ***, p = 0.0006 between ciliated wild type and G2019S mice; ns, no significance. Values represent the data (mean ± SEM) from individual brains, analyzing 4 brains per group, 2-3 sections per mouse, >30 neurons per mouse.

### Loss of PV neurons in the LRRK2 G2019S striatum

Kottman and colleagues have shown that blocking Shh expression in dopamine neurons of the substantia nigra leads to progressive degeneration of cholinergic and PV neurons in the mouse striatum (Gonzalez-Reyes et al., 2012). As shown in Figure 5, LRRK2 G2019S dorsal striatum lost almost 40% of PV neurons compared with wild type littermates, as detected by counting PV^+^ cell bodies across the entire dorsal striatum of four 5-month-old mutant mice. These data support the conclusion that pathogenic LRRK2 expression decreases the ability of striatal PV neurons to receive ciliary, neurotrophic Shh signals and to produce NRTN peptide in response. This phenotype occurs concomitant with decreased GDNF production by G2019S cholinergic neurons, exacerbating the loss of neuroprotection for dopamine neurons.

**Figure 5.**
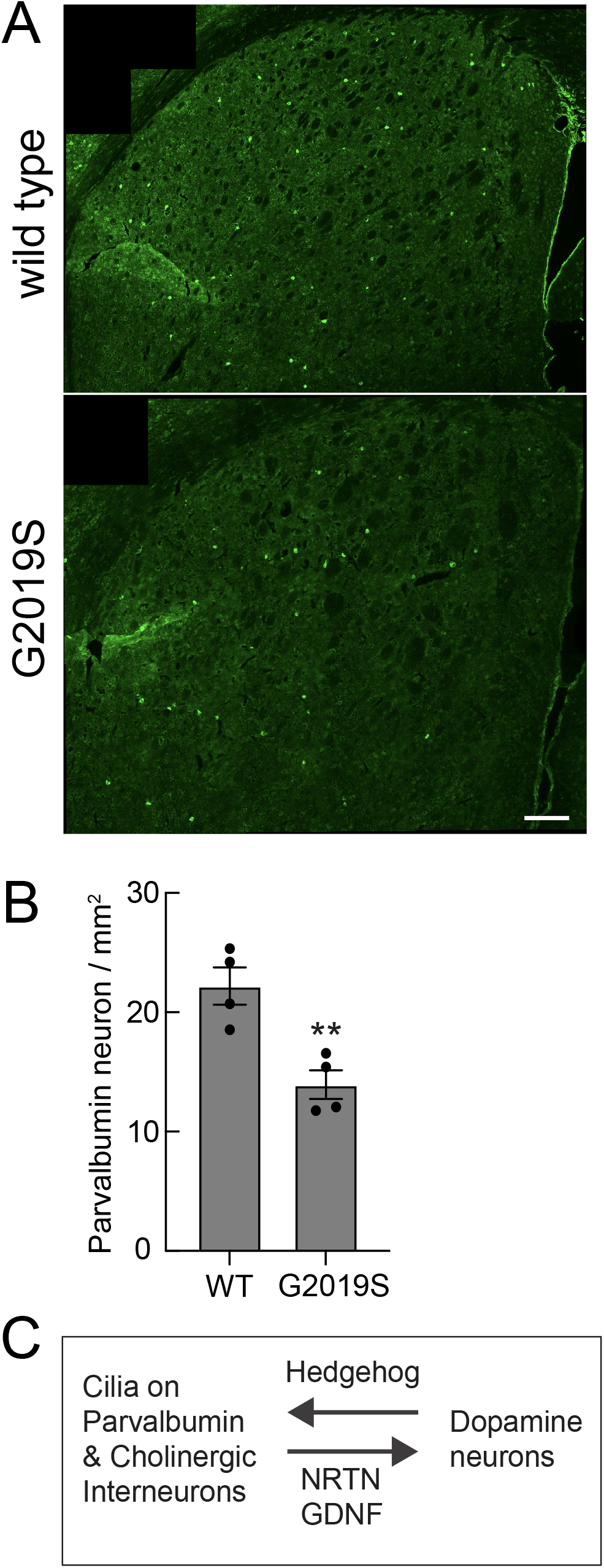
Loss of parvalbumin interneurons in G2019S LRRK2 dorsal striatum. (A) low magnification example images of total PV neurons from wild type (upper panel) and G2019S LRRK2 (lower panel) mice. Bar, 200 μm. (B) Quantitation of data from micrographs such as that shown in (A), detected per mm^2^. Significance was determined by unpaired *t*-test. **, p = 0.0058. Values represent the data (mean ± SEM) from individual brains, analyzing 4 brains per group, 2 sections per mouse, with >100 tiles scored per brain. (C). Reciprocal, cilia-dependent signaling between dopamine neurons of the Substantia nigra and striatal interneurons.

As summarized in Figure 5C, dopamine neurons infiltrate the striatum where they secrete Shh that is sensed by PV and cholinergic interneurons. Those cells rely on Hedgehog for viability (Gonzalez-Reyes et al., 2012) and provide neuroprotection to dopamine neurons by secreting NRTN and GDNF. In the absence of cilia, they cannot receive the Hedgehog signal and fail to produce NRTN and GDNF. They may also fail to survive in the absence of strong Hedgehog signals.

We have shown here that an activating mutation in LRRK2 kinase has profound effects on Shh signaling in PV neurons of the dorsal striatum. Cilia loss is accompanied by a significant decrease in PTCH1 expression in PV neurons, coupled with a loss of NRTN RNA. Indeed, both the decrease in PTCH1 and NRTN appear to be greater than the loss of overall ciliation in this cell type, but consistent with a combination of cilia loss and remaining ciliary shortening. Loss of Shh signaling leads to loss of PV neurons overall, consistent with prior studies showing their dependence on Shh for viability (Gonzalez-Reyes et al., 2012).

A remaining puzzle is why PV and cholinergic interneurons and striatal astrocytes are vulnerable to cilia loss whereas the surrounding, much more abundant medium spiny neurons are not, both in mice and in humans (Dhekne et al., 2018; Khan et al., 2024). It is not because they express higher levels of LRRK2; on the contrary, we have shown that they actually express less (Khan et al., 2024). Yet within the category of either PV interneurons (this study) or cholinergic interneuron cell types, higher expression correlates with decreased ciliogenesis. Further work will be needed to understand the unique vulnerability of interneuron- and astrocyte-cilia to LRRK2 kinase action.

The present study shows that Parkinson’s-associated LRRK2 mutation causes significant loss of NRTN expression in the mouse striatum due to loss of primary cilia in PV neurons that rely on cilia to produce NRTN in a Hedgehog responsive manner; GDNF expression in cholinergic interneurons is also decreased in these animals (Khan et al., 2024). Both GDNF and NRTN have been tested as therapeutics for PD patients as part of multiple clinical trials with mixed results to date (cf. (Barker et al., 2020; Chu and Kordower, 2023; Vastag, 2010)). A challenge for these trials has been to achieve adequate distribution of administered proteins or injected expression viruses, but alternative delivery strategies continue to be tried. These exogenous growth factors will hopefully help to sustain dopaminergic neurons. But without cilia to sense Shh, cholinergic and PV interneurons that are so important for regulating dopamine signaling will remain vulnerable and their numbers may nevertheless decrease (Gonzalez-Reyes et al., 2012). Altogether, these experiments highlight the pathogenic LRRK2-triggered loss of neuroprotection experienced by multiple neuron classes, due to a block in Shh signaling and neuroprotective factor production that has important implications for dopaminergic neuron survival in Parkinson’s disease.

## Materials and Methods

### Research standards for animal studies

Mice were obtained from Taconic (Constitutive KI Lrrk2tm4.1Arte; RRID:I MSR_TAC:13940) and kept in specific pathogen-free conditions at the University of Dundee (UK). All animal experiments were ethically reviewed and conducted in compliance with the Animals (Scientific Procedures) Act 1986 and guidelines established by the University of Dundee and the U.K. Home Office. Ethical approval for animal studies and breeding was obtained from the University of Dundee ethical committee, and all procedures were performed under a U.K. Home Office project license. The mice were group-housed in an environment with controlled ambient temperature (20–24°C) and humidity (45–55%), following a 12-hour light/12-hour dark cycle, with ad libitum access to SDS RM No. three autoclavable food and water. Genotyping of mice was performed by PCR using genomic DNA isolated from ear biopsies. Primer 1 (5’ - CTGCAGGCTACTAGATGGTCAAGGT – 3’) and Primer 2 (5’ – CTAGATAGGACCGAGTGTCGCAGAG- 3’) were used to detect the wild-type and knock-in alleles (dx.doi.org/10.17504/protocols.io.5qpvo3xobv4o/v1). Homozygous LRRK2-G2019S mice and their littermate wild-type controls (5 months old) were utilized for the experiments, with genotyping confirmation conducted on the day of experiment.

### Immunohistochemical (IHC) staining

The mouse brain striatum was subjected to immunostaining following a previously established protocol (dx.doi.org/10.17504/protocols.io.bnwimfce). Frozen slides were thawed at room temperature for 15 minutes and then gently washed twice with PBS for 5 minutes each. Antigen retrieval was achieved by incubating the slides in 10 mM sodium citrate buffer pH 6.0, preheated to 95°C, for 15 minutes. Sections were permeabilized with 0.1% Triton X-100 in PBS at room temperature for 15 minutes, followed by blocking with 2% FBS and 1% BSA in PBS for 2 hours at room temperature. Primary antibodies were applied overnight at 4°C, and the next day, sections were exposed to secondary antibodies at room temperature for 2 hours. Secondary antibodies used were donkey highly cross-absorbed H + L antibodies conjugated to Alexa or CF™488, Alexa 568, or Alexa 647, diluted at 1:2000. Nuclei were counterstained with 0.1 μg/ml DAPI (Sigma). Finally, stained tissues were mounted with Fluoromount G and covered with a glass coverslip. All antibody dilutions for tissue staining contained 1% DMSO to facilitate antibody penetration.

### Fluorescence in situ hybridization (FISH)

RNAscope fluorescence in situ hybridization was carried out as described (https://bio-protocol.org/exchange/preprintdetail?id=1423&type=3&searchid=EM1708992000021453&sort=5&pos=b;(Khan et al., 2021)). The RNAscope Multiplex Fluorescent Detection Kit v2 (Advanced Cell Diagnostics) was utilized following the manufacturer’s instructions, employing RNAscope 3-plex Negative Control Probe (#320871) or Mm-Lrrk2 (#421551), Mm-Ptch1-C2 (#402811-C2) and Mm-Nrtn-C2 (#441501-C2). The Mm-Lrrk2, Mm-Ptch1-C2 and Mm-Nrtn-C2 probes were diluted 1:20, 1:5 and 1:3, respectively in dilution buffer consisting of 6x saline-sodium citrate buffer (SSC), 0.2% lithium dodecylsulfate, and 20% Calbiochem OmniPur Formamide. Fluorescent visualization of hybridized probes was achieved using Opal 690 (Akoya Biosciences). Subsequently, brain slices were subjected to blocking with 1% BSA and 2% FBS in TBS (Tris buffered saline) with 0.1% Triton X-100 for 30 minutes. They were then exposed to primary antibodies overnight at 4°C in TBS supplemented with 1% BSA and 1% DMSO. Secondary antibody treatment followed, diluted in TBS with 1% BSA and 1% DMSO containing 0.1 μg/ml DAPI (Sigma) for 2 hours at room temperature. Finally, sections were mounted with Fluoromount G and covered with glass coverslips.

### Microscope image acquisition

All images were obtained using a Zeiss LSM 900 confocal microscope (Axio Observer Z1/7) coupled with an Axiocam 705 camera and immersion objective (Plan-Apochromat 63x/1.4 Oil DIC M27) or objectives (Plan-Apochromat 20x/0.8 M27 and EC Plan Neofluar 5x/0.16 M27). The images were acquired using ZEN 3.4 (blue edition) software, and visualizations and analyses were performed using Fiji.

In addition to above-mentioned methods, all other statistical analysis was carried out using GraphPad Prism version 9.3.1 for Macintosh, GraphPad Software, Boston, Massachusetts USA, www.graphpad.com.

## Acknowledgments

This study was funded by the joint efforts of The Michael J. Fox Foundation for Parkinson’s Research (MJFF) and Aligning Science Across Parkinson’s (ASAP) initiative. MJFF administers the grant (ASAP-000463) on behalf of ASAP and itself. For the purpose of open access, the authors have applied a CC-BY public copyright license to the Author Accepted Manuscript version arising from this submission.

## Author contributions

YL and EJ, Conceptualization, investigation, formal analysis, data curation, visualization, manuscript editing; FT, Project administration, resources; SRP, Conceptualization, writing original draft, supervision, formal analysis, funding acquisition, visualization.

The authors declare that they have no competing interests. All primary data is available at: https://doi.org/10.5281/zenodo.11510060

## Figure Legends

**Supplemental Table 1.** Key Resources used in this study.

| Reagent type (species) or resource | Designation | Source or reference | Identifiers | Additional information |
| --- | --- | --- | --- | --- |
| Genetic reagent(Mus musculus) | Constitutive KI Lrrk2tm4.1Arte | Taconic | #13940,<br>RRID:IMSR_TAC:13940 | C57BL/6;<br>G2019S KI |
| Antibody | anti-Parvalbumin (guinea pig polyclonal) | Synaptic system | #195004<br>(RRID:AB_2156476) | (1:1000) |
| Antibody | anti-Somatostatin 28 (guinea pig polyclonal) | Synaptic system | #366004<br>(RRID:AB_2620126) | (1:500) |
| Antibody | anti-Adenylate cyclase III (rabbit polyclonal) | EnCOR | RPCA-ACIII<br>(RRID:AB_2572219) | (1:10000) |
| Antibody | H+L Donkey anti-guinea pig Alexa 647 | Jackson ImmunoResearch | #706-605-148<br>(RRID:AB_2340476) | (1:2000) |
| Antibody | H+L Donkey anti-guinea pig CF<sup>TM</sup> 488A | Sigma-Aldrich | SAB4600033<br>(RRID:AB_2890881) | (1:2000) |
| Antibody | H+L Donkey anti-Rabbit Alexa 568 | Life Technologies | A10042<br>(RRID:AB_2534017) | (1:2000) |
| Commercial assay or kit | RNAscopeMultiplex Fluorescent Reagent Kit v2 | Advanced Cell Diagnostics | #323100 |  |
| Commercial assay or kit | RNAscope Probe-Mm-Lrrk2 | Advanced Cell Diagnostics | #421551 | (1:20) |
| Commercial assay or kit | RNAscope Probe- Mm-Ptch1-C2 | Advanced Cell Diagnostics | #402811-C2 | (1:5) |
| Commercial assay or kit | RNAscope Probe- Mm-Nrtn-C2 | Advanced Cell Diagnostics | #441501-C2 | (1:3) |
| Commercial assay or kit | OPAL 690 REAGENT PACK | Akoya Biosciences | FP1497001KT |  |
| Software, Algorithm | FIJI | PMID:29187165 | RRID:SCR_002285 |  |
| Software, Algorithm | Prism | Prism 9 version 9.3.1 | RRID:SCR_002798 |  |
| Software, Algorithm | ZEN | ZEISS ZEN Microscopy Software | RRID:SCR_013672<br>www.zeiss.com/microscopy/en/products/software/zeiss-zen.html |  |

## Abbreviations

ACIII: Adenylate cylase 3
GABA: 
GDNF: glial-derived neurotrophic factor
LRRK2: Leucine-rich repeat kinase 2
NRTN: Neurturin
PTCH1: Patched 1
PV: Parvalbumin
SST28: Somatostatin Receptor 28

